# Human basal-like breast cancer is represented by one of the two mammary tumor subtypes in dogs

**DOI:** 10.1101/2023.03.02.530622

**Authors:** Joshua Watson, Tianfang Wang, Kun-Lin Ho, Yuan Feng, Kevin K Dobbin, Shaying Zhao

## Abstract

**Background:** About 20% of breast cancers in humans are basal-like, a subtype that is often triple negative and difficult to treat. An effective translational model for basal-like breast cancer (BLBC) is currently lacking and urgently needed. To determine if spontaneous mammary tumors in pet dogs could meet this need, we subtyped canine mammary tumors and evaluated the dog-human molecular homology at the subtype level.

**Methods:** We subtyped 236 canine mammary tumors from 3 studies by applying various subtyping strategies on their RNA-seq data. We then performed PAM50 classification with canine tumors alone, as well as with canine tumors combined with human breast tumors. We investigated differential gene expression, signature gene set enrichment, expression association, mutational landscape, and other features for dog-human subtype comparison.

**Results:** Our independent genome-wide subtyping consistently identified two molecularly distinct subtypes among the canine tumors. One subtype is mostly basal-like and clusters with human BLBC in cross-species PAM50 classification, while the other subtype does not cluster with any human breast cancer subtype. Furthermore, the canine basal-like subtype recaptures key molecular features (e.g., cell cycle gene upregulation, TP53 mutation) and gene expression patterns that characterize human BLBC. It is enriched histological subtypes that match human breast cancer, unlike the other canine subtype. However, about 33% of canine basal-like tumors are estrogen receptor negative (ER-) and progesterone receptor positive (PR+), which is rare in human breast cancer. Further analysis reveals that these ER-PR+ canine tumors harbor additional basal-like features, including upregulation of genes of interferon-γ response and of the Wnt-pluripotency pathway. Interestingly, we observed an association of *PGR* expression with gene silencing in all canine tumors, and with the expression of T cell exhaustion markers (e.g., *PDCD1*) in ER-PR+ canine tumors.

**Conclusions:** We identify a canine mammary tumor subtype that molecularly resembles human BLBC overall, and thus could serve as a vital spontaneous animal model of this devastating breast cancer subtype. Our study also sheds light on the dog-human difference in the mammary tumor histology and the hormonal cycle.

## Background

Human breast cancer is heterogeneous, consisting of well-established molecularly distinct subtypes [1–10]. One of these subtypes is basal-like breast cancer (BLBC; human BLBC will be referred to as hBLBC hereafter), which makes up roughly 15-20% of human breast cancers and has the worst prognosis of all subtypes [1–10]. About 70% of hBLBCs are triple negative, expressing neither estrogen receptor (ER) nor progesterone receptor (PR) and without HER2 amplification or overexpression [1–10]. These cancers also tend to have increased rates of cell proliferation and metastasis [1–10]. All these highlight the need for an effective translational model for hBLBC, which is critically missing at present [11–13].

Mammary cancers in pet dogs naturally occur in animals with an intact immune system [13, 14], overcoming many limitations of traditional cancer models such as cell lines and genetically modified rodent models. These canine cancers more accurately emulate human breast cancers in etiology, complexity, heterogeneity, behavior, treatment, and outcome [13–16]. They are also common in bitches, with an annual incidence rate estimated at 198 per 100,000 [17], which is comparable to the rate of 125 per 100,000 for breast cancer in women in the United States [18]. Mammary cancer is especially common in bitches that are not spayed or are spayed after the second estrus, with the risk for malignant tumor development expected at 26% [17]. Thus, canine mammary tumors have the potential to serve as a much-needed translational model of hBLBC, effectively bridging a current gap between preclinical models and human clinical trials to accelerate bench-to-bedside translation.

The effective use of the canine model is, however, complicated by issues including the dog-human hormonal cycle difference, e.g., the luteal phase lasts ∼14 days for humans but ∼2 months for dogs. Another difference is histology. About 50% of canine mammary tumors are complex or mixed, with multiple cell lineages (e.g., epithelial and myoepithelial cells) proliferating [14, 19, 20]. These histologies (e.g., adenomyoepithelioma) are, however, are very rare (<1%) in human breast cancers [14, 21, 22]. It remains unknown how these differences shape the molecular homology and difference between canine and human mammary tumors.

The same as human breast cancers, spontaneous canine mammary cancers are heterogeneous and consist of distinct subtypes [19, 20]. Thus, subtype level dog-human comparison is needed to evaluate the dog-human homology. Canine mammary cancers have been histopathologically and clinically subtyped, as well as molecularly subtyped with immunohistochemical markers established for human breast cancer (anti-ER, PR, HER2, -CK 5/6 and -CK14) [19, 20, 23, 24]. However, to our knowledge, canine mammary tumors have not been independently subtyped by genome-wide molecular studies.

For dog-human comparison, our previous study indicates that one histological subtype, simple carcinoma, molecularly resembles hBLBC in cross-species PAM50 classification with human and canine tumors [14]. However, this study is limited by its small sample size. Another group has performed PAM50 classification on the RNA-seq data recently published for 154 canine mammary tumors [25], and reported a higher homology between canine and human luminal A tumors than between canine and human basal-like tumors [26]. This study, however, did not perform cross-species PAM50 to directly compare human and canine tumors. Moreover, many of the canine luminal A tumors are complex and mixed tumors, histologically differing from the vast majority of human luminal A tumors [14, 21, 22].

To address these discrepancies and deficiencies, we set out to independently subtype canine mammary tumors using RNA-seq data of 236 tumors [14, 25, 27], and then perform dog-human comparison at the subtype level, as described below.

## Materials and Methods

### Data collection

Canine RNA-seq data was downloaded from the Sequence Read Archive (SRA) database, including data of 154 mammary tumors from PRJNA489087 (excluding 4 metastatic osteosarcoma and fibrosarcomas) [25] and of 63 mammary tumors from PRJNA561580 [27]. RNA-seq data of 25 mammary tumor samples sequenced in house [14] were also included (PRJNA203086 and PRJNA912710). Human breast cancer RNA-seq data were downloaded from the National Cancer Institute (NCI) Genomic Data Commons (GDC) database, and the PAM50 classification of these cancers was obtained from the cBioportal database [28]. Gene expression microarray data of two canine mammary tumor studies (GSE20718 and GSE22516) and one human breast cancer (GSE20685) were downloaded from the Gene Expression Omnibus (GEO) database [29–31], and processed with affyPackage [32]. Other information was obtained from relevant publications of these studies. Canine genome canFam3.1 and gene annotation canFam3 1.99 GTF were downloaded from the Ensembl database. Canine mutation data and tumor mutation burden (TMB) values from whole exome sequencing analysis were obtained from a previous publication [33].

### Canine sample collection and RNA-seq

Fresh-frozen (FF) canine tissues and spontaneous tumors were obtained from the canine tissue archive bank at Ohio State University, as previously described [34]. Samples were collected from client-owned dogs that developed the disease spontaneously, under the guidelines of the Institutional Animal Care and Use Committee for use of residual diagnostic specimens and with owner informed consent. The case information was provided by the tissue bank. The research received the ethical approval from the Institutional Animal Care and Use Committee.

Cryosectioning of FF tissues, H&E staining, and cryomicrodissection were performed as described [34] to enrich tumor cells for tumor samples. Genomic DNA and RNA were extracted from the dissected tissues using the AllPrep DNA/RNA Mini Kit (cat. no. 80204) from QIAGEN (Germantown, MD, USA). Only samples with a 260/280 ratio of ∼2.0 (RNA) and showing no degradation and other contaminations were subjected to further quality control with qRT-PCR analysis with a panel of genes as previously described [34, 35]. RNA-seq libraries were constructed using KAPA Stranded mRNA-Seq Kit. The samples were subjected to 75 or 125 bp paired-end sequencing using Illumina HiSeq 2500 or NextSeq 500 at Georgia Genomics Facility.

### Canine RNA-seq data quality control (QC) and processing

Canine RNA-seq data were processed as described [34–36]. Briefly, RNA-seq read pairs were mapped to the canine reference genome canFam3 using HISAT2 (version 2.21) [37]. Concordantly (for paired-end RNA-seq data only) and uniquely mapped pairs were identified and were used to calculate the mapping rate of each sample. Such pairs with at least one read with ≥1 bp overlapping a coding sequence (CDS) region of the canFam3 1.99 GTF annotation were used to calculate the CDS-targeting rate.

Quality control of canine RNA-seq data was performed as described [36]. First, MultiQC [38] (version 1.5) was used to examine GC content and duplicate level. Base quality distribution before and after Trimmomatic trimming was also examined. Second, the distributions of per sample read-pair total amount, mapping quality, and CDS targeting rate were examined to identify and exclude samples that fail to meet the cutoffs. A total of 6 canine RNA-seq samples from PRJNA561580 failed the QC and were excluded from further analysis.

For each sample that passed QC measures, Subread (version 2.0.0) [39] was used to identify read pairs that are uniquely and, for paired-end RNA-seq, concordantly mapped to the exonic regions of the canFam3 1.99 GTF annotation, the sum of which yields raw RNA-seq counts. Cufflinks version 2.2.0 [40] was used to calculate the FPKM (fragments per kilobase of exon per million mapped) value of each gene in each sample, which was then converted to TPM (transcript per million). For studies that combine RNA-seq data from multiple sources, comBat [41] was applied to correct batch effect for TPM values, and ComBat-seq [42] was used to correct batch effect for mapped RNA-seq read count values.

### ER, PR, and HER2 status

Samples with an *ESR1* or *PGR* expression level of 𝐹𝑃𝐾𝑀 ≤ 1 and 𝐹𝑃𝐾𝑀 > 1 were classified as ER or PR negative and positive respectively. Samples with an *ERBB2* expression level of 𝐹𝑃𝐾𝑀 ≤ 35 and 𝐹𝑃𝐾𝑀 > 35 were classified as HER2 not enriched and enriched, respectively.

### Canine mammary tumor subtyping

A gene was selected if it has an official gene name associated with its Ensembl gene ID in the canFam3 1.99 GTF annotation and is expressed (with 𝐹𝑃𝐾𝑀 ≥ 1 in at least one sample across a cohort or study). This yields 13,416 genes in the discovery set (paired-end RNA-seq data) [14, 25] and 13,608 genes in the validation set (single-end RNA-seq data) [27] (see Results). The NMF R package [43] was then applied on all of these selected genes, as well as on the top 5000, 2000, 1000, and 500 most variable genes among them, with 30 runs for the rank determination. These analyses consistently divided 143 out of 179 samples of the discovery set into two subtypes. For validation, K-means clustering, consensus clustering, and hierarchical clustering via R packages stats, ConsensusClusterPlus, and pvclust respectively [44–46] were used to subtype the 143 samples using the top 10% most variable genes. The same process was repeated to subtype the samples in the validation set.

### PAM50 classification of canine and human tumors

A total of 43 canine homologues of the 50 PAM50 genes were identified in the canFam3 1.99 annotation file. These 43 genes were then used to perform PAM50 classification with canine samples alone, as well as with canine and human combined samples, for RNA-seq studies [14, 25, 27]. For microarray studies [30, 31], 40 of the PAM50 genes were identified based on the probes and used to classify canine samples alone and canine-human combined samples. The PAM50 subtypes of human breast cancer samples were downloaded from the cBioportal database or relevant publications. For canine and human combined sample PAM50 clustering for RNA-seq studies, 60 human tumors were randomly sampled from the cancer genome atlas (TCGA) breast cancer study for each of the luminal A, luminal B, hBLBC, and HER2-enriched subtypes. These samples, along with all 27 human normal-like tumors, were merged with all 143 subtyped canine tumors for PAM50 classification analysis. The clustering dendrogram was then cut at the minimum number of clusters that maximally separate hBLBC tumors from hLumA tumors using the R package dendextend [47]. The number of tumors of each canine or human subtype in each cluster was counted. If a cluster contains the majority of tumors of a human subtype as well as the majority of a canine subtype, the human and canine subtypes were considered matched. This process was repeated 100 times to ensure each human tumor was sampled at least once.

Multidimensional scaling on the Euclidean distance matrix was performed for each of the 100 random samplings, from which the mahalanobis distance between the centers of any two subtypes was calculated using the R package ‘GenAlgo’ v2.2.0 [48].

The above process was repeated for microarray studies [29–31], except that only 40 canine homologues of the 50 PAM50 genes were identified, and 30 tumors per subtype were randomly sampled from the human dataset [29].

### Differentially expressed (DE) genes and gene set enrichment analysis (GSEA)

DESeq2 [49] was used to identify DE genes between subtypes or subgroups. Genes with ≥ 2 fold change in read count and the Benjamini-Hochberg (BH) adjusted 𝑝 ≤ 0.05 were considered differentially expressed. Enriched functions of DE genes were investigated with the GSEA [50] and DAVID [51] web tools. Pathway and signature gene sets were acquired from previous publications [34, 35, 52]. Single sample GSEA (ssGSEA v. 10.1.0) was performed using Genepattern [53].

### Correlation and other statistical analyses

Genes with 𝐹𝑃𝐾𝑀 > 1 in at least 10% of the samples of a subtype were chosen for *ESR1* or *PGR* correlation analysis. *PGR* was excluded from hBLBC as <10% of samples have an 𝐹𝑃𝐾𝑀 > 1. Positively or negatively correlated genes were defined as those with BH adjusted 𝑝 ≤ 0.05 and correlation coefficient |𝑅| ≥ 0.3 for both Pearson and Spearman correlation analysis. The software enrichR [54] was used to identify transcription factors targeting each group of significantly correlated genes. Wilcoxon rank sum tests were used for statistical comparison between subtypes or subgroups.

## Results

### Canine mammary tumors consist of two distinct subtypes

We first performed non-negative matrix factorization (NMF) [43], a widely used subtyping strategy, to subtype 179 canine mammary tumors with paired-end RNA-seq data (the discovery set), after combining 154 tumors sequenced by Kim et al. [25] and 25 tumors sequenced by us [14] followed by batch correction. NMF subtyping was repeated with all 13,416 genes that are expressed in at least one tumor, as well as with the top 5000, 2000, 1000, and 500 most variable genes among the 13,416 expressed genes. The analysis consistently clustered 143 of 179 tumors into two subtypes (Fig. 1). To validate this, we also subtyped these tumors using other popular strategies, including K-means, consensus clustering, and hierarchical clustering via multiscale bootstrap resampling [44, 45]. These strategies consistently identified the same two subtypes as the NMF approach.

**Fig. 1.**
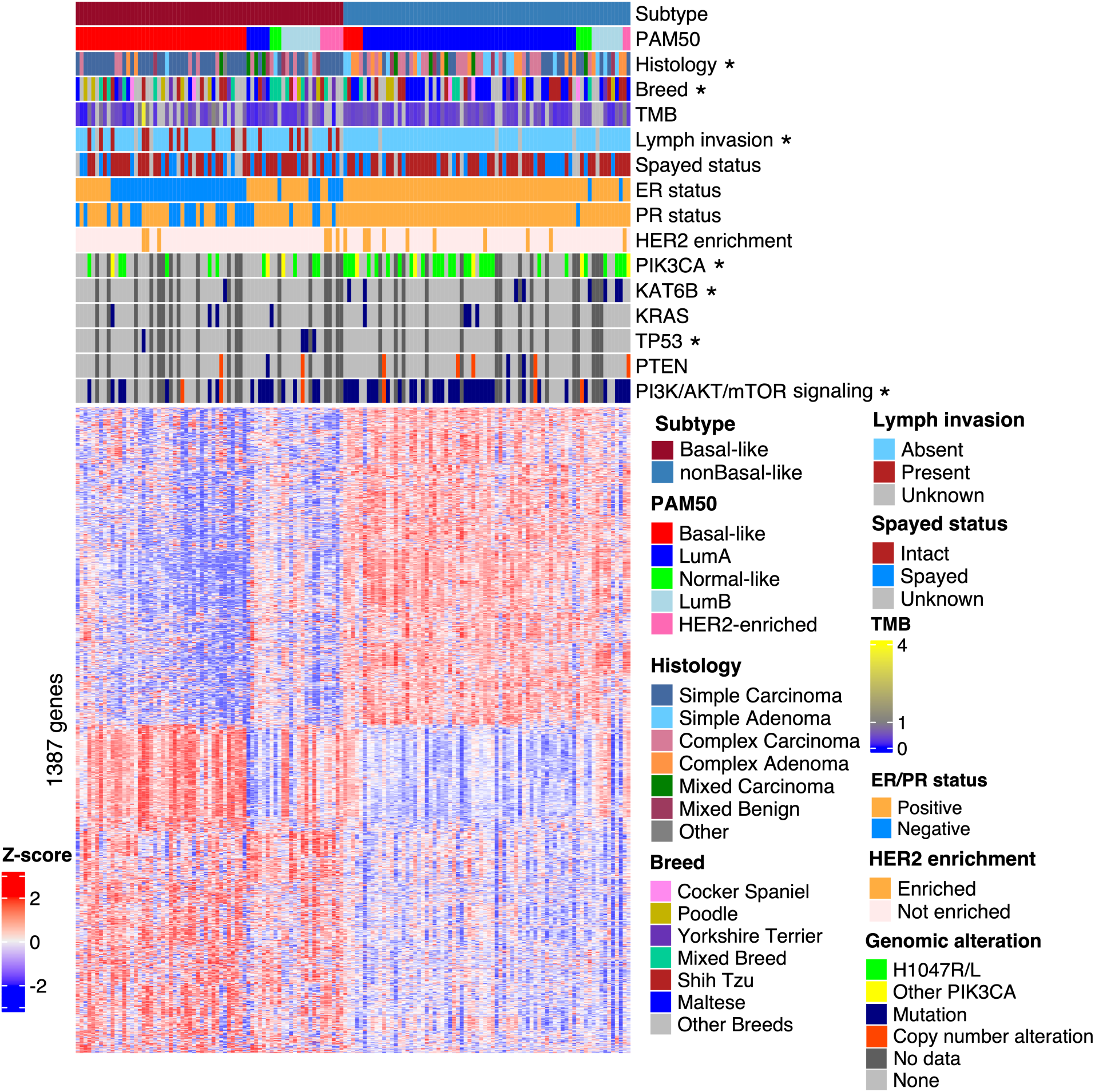
Two molecularly distinct subtypes of canine mammary tumors were identified. Heatmap of the row scaled log_2_(TPM) values of 1,387 metagenes from 143 canine mammary tumors (columns), ordered from left to right by subtypes, PAM50 classification, and the *ESR1* expression level from high to low. The metagenes (rows), identified from the NMF analysis (see Methods), are ordered by hierarchical clustering. Lymph invasion is defined as having tumor cells in peritumoral lymphatic vessels and/or regional lymph nodes. Tumor mutation burden (TMB), defined as the number of somatic base substitutions and small indels per megabase (Mb) of callable coding sequence, is obtained from a previous publication [33]. For ER/PR status, a tumor is considered “negative” or “positive” if its *ESR1*/*PGR* has a FPKM value of ≤ 1 or > 1, respectively. For HER2 enrichment, a sample is considered “not enriched” or “enriched” if its *ERBB2* has a FPKM value of ≤ 35 or >35, respectively. “Other PIK3CA” represents all non-H1047 coding mutations in PIK3CA. “No data” represents samples with no mutation data. Annotation row titles marked with a “*” indicate a significant (𝑝 ≤ 0.05) difference in enrichment between the subtypes.

We then performed the same subtyping analyses on the other large study of canine mammary tumors (n=57) (the validation set), whose RNA-seq data are single-end [27], differing from the discovery set. These tumors also clustered into two subtypes, consistent with the discovery set.

The two subtypes differ in tumor histology and invasiveness. One subtype is significantly (𝑝 = 0.007) enriched in simple adenomas/carcinomas (where only one cell lineage proliferates prominently), while the other subtype is enriched in complex or mixed adenomas/carcinomas (𝑝 < 0.001) (where more than one cell lineages are proliferating prominently) (Fig. 1) [14, 19, 25]. Moreover, the simple adenomas/carcinomas-enriched subtype contains significantly (𝑝 < 10^−6^) more cases with lymph node invasion (Fig. 1). Interestingly, the other subtype contains significantly (𝑝 = 0.0014) more Maltese dogs (Fig. 1).

The two subtypes display distinct molecular features. Among the top most mutated genes in this cohort [33], one subtype is enriched in *PIK3CA* hotspot mutation H1047R/L (𝑝 = 0.0034) and *KAT6B* mutation (𝑝 = 0.036), while the other subtype (simple adenomas/carcinomas-enriched and with more lymph node invasion) is enriched in *TP53* mutation (𝑝 = 0.043) (Fig. 1). However, we did not observe any significant difference in *KRAS* mutation and TMB between the two subtypes (Fig. 1). For pathways, the two subtypes differ in mutations and copy number alterations of genes in PI3K signaling (𝑝 = 0.0039) (Fig. 1), the most altered pathway in canine mammary tumors [33].

Other molecular differences between the two subtypes include PAM50 classification, ER and PR expression status, and others, and will be described in more details below.

### Canine and human basal-like tumors cluster together in PAM50 classification

We performed PAM50 classification [55, 56] on the 179 canine tumors from the discovery set. About 74% of the tumors were classified as either the basal-like (n=62, 35%) or luminal A (n=71, 39%) subtype (Fig. 2A; Table 1). Importantly, 90% of the basal-like tumors belong to one of the two subtypes shown in Fig. 1, while 90% of the luminal A tumors belong to the other subtype. For this reason and reasons described below, the two subtypes shown in Fig. 1 are named canine basal-like mammary tumor (cBLMT) and canine non-basal-like mammary tumor (cNBLMT) respectively.

**Fig. 2.**
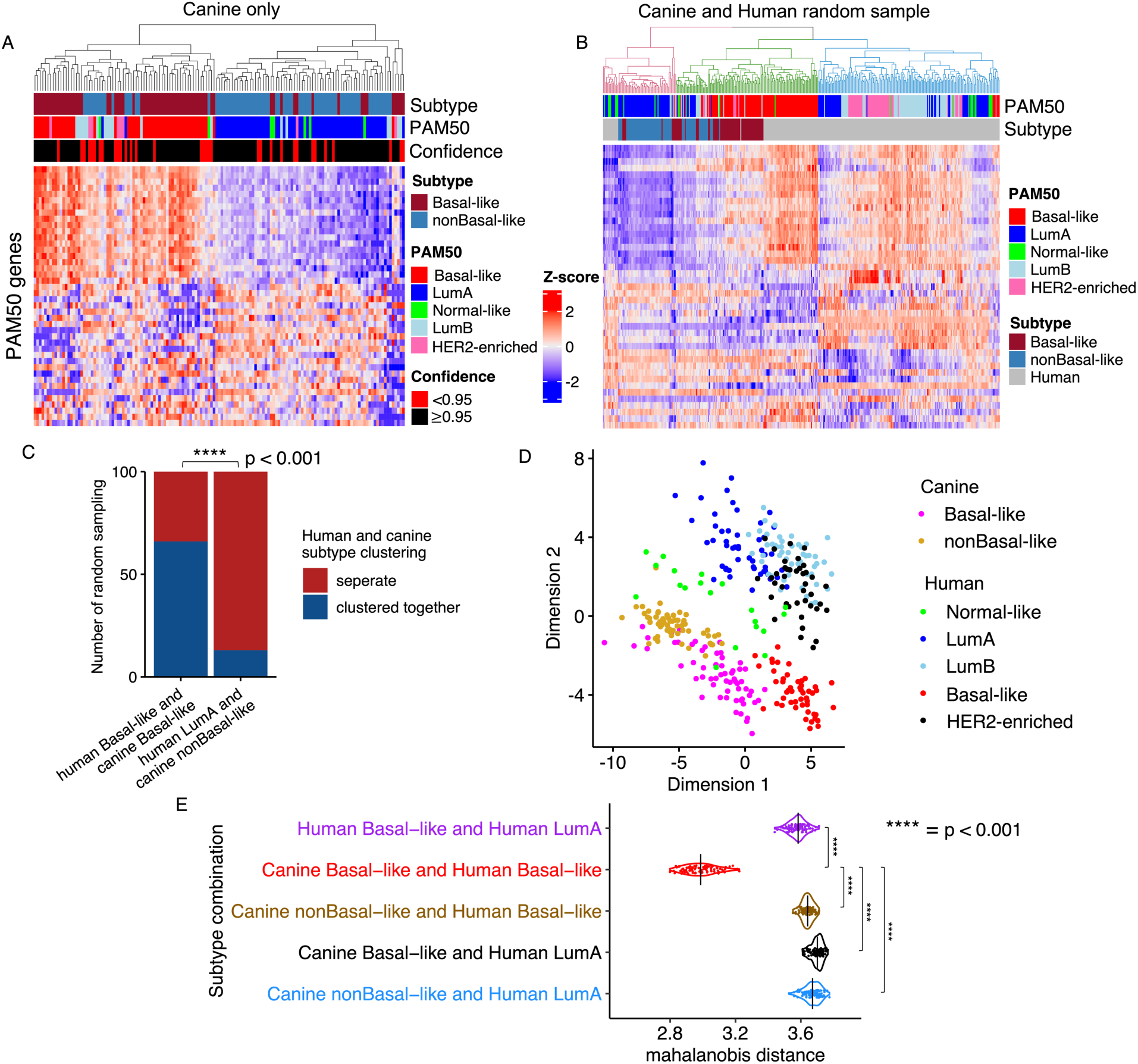
PAM50 classification groups canine and human basal-like tumors, but separates canine non-basal-like and human tumors. A. PAM50 classification of 143 subtyped canine mammary tumors. The heatmap shows hierarchical clustering of the canine tumors using the row scaled log_2_(TPM) values of 43 out of the 50 PAM50 genes. The bars indicate the canine subtype from Fig. 1, as well as the PAM50 subtype and confidence score of each tumor. B. An example of cross-species PAM50 classification. All 143 subtyped canine tumors, together with 267 human tumors (60 tumors per subtype of luminal A (LumA), luminal B (LumB), basal-like or HER2-enriched randomly sampled from TCGA database, along with all 27 normal-like tumors from TCGA) were subjected to PAM50 classification. The dendrogram is colored to indicate the minimum clusters that maximally separate the hLumA tumors from hBLBC tumors. This cross-species PAM50 classification were repeated 100 times. C. Bar plot showing the number of random samplings in which human and canine basal-like tumors clustered together or separately, compared to those of human luminal A and canine non-basal-like tumors. The p-value is based on Fisher’s exact test. D. Multidimensional scaling plot of the cross-species PAM50 classification shown in B. Each dot represents a tumor from a subtype specified by the color as indicated in the legend. E. Violin plot indicating the distribution of the mahalanobis distances between the centers of two subtypes on the multidimensional scaled plot (D) of each of the 100 cross-species PAM50 classifications (B) achieved via random samplings (see Methods). The p-values were obtained from Wilcoxon tests

**Table 1.**
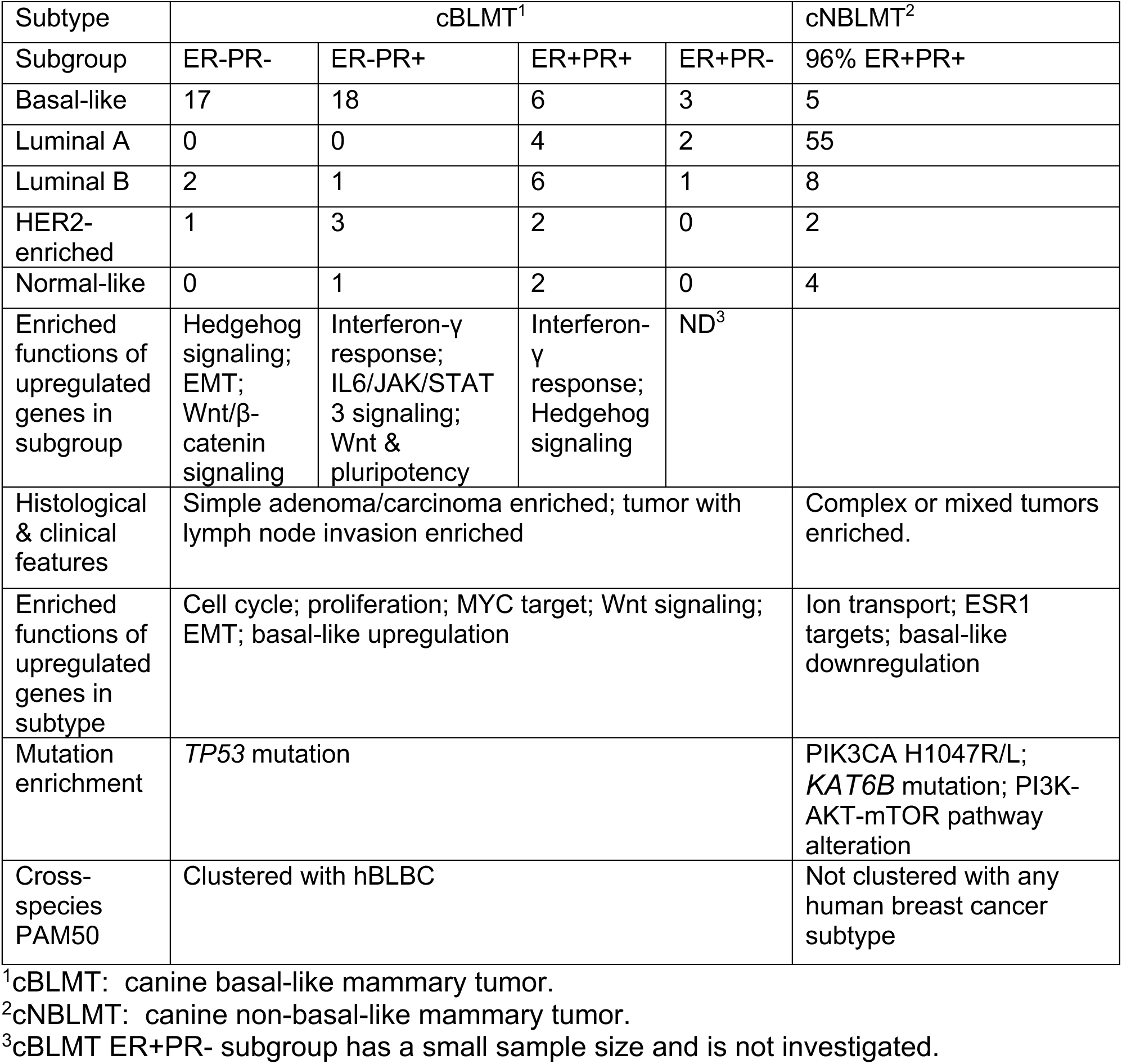
Features of canine mammary tumor subtypes and subgroups

| Subtype | cBLMT<sup>1</sup> |  |  |  | cNBLMT<sup>2</sup> |
| --- | --- | --- | --- | --- | --- |
| Subgroup | ER-PR- | ER-PR+ | ER+PR+ | ER+PR- | 96% ER+PR+ |
| Basal-like | 17 | 18 | 6 | 3 | 5 |
| Luminal A | 0 | 0 | 4 | 2 | 55 |
| Luminal B | 2 | 1 | 6 | 1 | 8 |
| HER2-enriched | 1 | 3 | 2 | 0 | 2 |
| Normal-like | 0 | 1 | 2 | 0 | 4 |
| Enriched functions of upregulated genes in subgroup | Hedgehog signaling; EMT; Wnt/β-catenin signaling | Interferon-γ response; IL6/JAK/STAT 3 signaling; Wnt & pluripotency | Interferon-γ response; Hedgehog signaling | ND<sup>3</sup> | |
| Histological & clinical features | Simple adenoma/carcinoma enriched; tumor with lymph node invasion enriched |  |  |  | Complex or mixed tumors enriched. |
| Enriched functions of upregulated genes in subtype | Cell cycle; proliferation; MYC target; Wnt signaling; EMT; basal-like upregulation |  |  |  | Ion transport; ESR1 targets; basal-like downregulation |
| Mutation enrichment | *TP53* mutation |  |  |  | PIK3CA H1047R/L; *KAT6B* mutation; PI3K-AKT-mTOR pathway alteration |
| Cross-species PAM50 | Clustered with hBLBC |  |  |  | Not clustered with any human breast cancer subtype |

<sup>1</sup>cBLMT: canine basal-like mammary tumor.
<sup>2</sup>cNBLMT: canine non-basal-like mammary tumor.
<sup>3</sup>cBLMT ER+PR- subgroup has a small sample size and is not investigated.

To quantitatively assess the canine-human homology at the subtype level, we performed cross-species PAM50 classification as described [14]. Briefly, we randomly sampled 60 human tumors from each of the luminal A, luminal B, HER2-enriched, and basal-like subtypes from the TCGA RNA-seq study [2, 3]. These, along with all 27 normal-like tumors in TCGA, amount to 267 human tumors covering all five intrinsic subtypes. We then performed PAM50 clustering on these human tumors together with all 143 subtyped canine tumors shown in Fig. 1. This analysis was repeated 100 times, ensuring that each TCGA tumor was sampled at least once.

In 66 of 100 random sampling analyses, cBLMTs and hBLBCs clustered together and away from tumors of any other canine or human subtype (Figs. 2B-C). To the contrary, in 87 of 100 random sampling analyses, cNBLMTs, the other canine subtype, did not cluster with tumors of any human subtype (Figs. 2B-C). These observations are supported by multidimensional scaling of each cross-species PAM50 classification, as shown by the example provided in Fig. 2D. We then calculated the mahalanobis distance [48] between the centers of canine and/or human subtypes for each of the 100 random sampling analyses. The distributions clearly indicate that the mahalanobis distances between hBLBC and cBLMT are significantly shorter than those between hBLBC and human luminal A (hLumA) or cNBLMT, as well as those between hLumA and cBLMT or cNBLMT (Fig. 2E). These results support that cBLMT molecularly resembles hBLBC, but cNBLMT molecularly differs from hLumA.

To validate this finding, we attempted to conduct the same analyses on the 57 tumors from the validation set [27]. PAM50 analysis of these canine tumors alone indeed classified a majority of the tumors as basal-like (n=21) or luminal A (n=17), consistent with the discovery set (Fig. 2A). Moreover, many of the basal-like canine tumors have a PAM50 gene expression patten that closely matches hBLBC. However, likely due to having single-end RNA-seq data, these canine tumors could not co-cluster with human breast tumors (whose RNA-seq data are paired-end) in cross-species PAM50 classification even after batch correction.

We next performed the same analyses on the gene expression microarray data of two canine studies (n=27; n=13) and of one human study (n=327) [29–31]. Consistent with the RNA-seq analysis described above, canine-only PAM50 clustering classified most of these canine tumors as either basal-like or luminal A. Moreover, cross-species PAM50 classification clustered cBLMTs and hBLBCs together in a majority of random sampling analyses, further supporting the molecular homology between cBLMT and hBLBC.

### The cBLMT subtype captures key molecular features of hBLBC

We identified differentially expressed (DE) genes between cBLMT and cNBLMT. The 1,123 genes upregulated in cBLMT are significantly enriched in cell cycle (e.g., DREAM targets, G2M checkpoint) and other functions that characterize hBLBC [3], as well as in genes that are known to be upregulated in hBLBC [52] (Fig. 3A; Table 1). Conversely, the 497 genes downregulated in cBLMT are significantly enriched in genes that are known to be downregulated in hBLBC [52], as well as in functions including ion transport and *ESR1* targets (Fig. 3A; Table 1). We conducted the same analysis with the validation set, and observed similar findings.

**Fig. 3.**
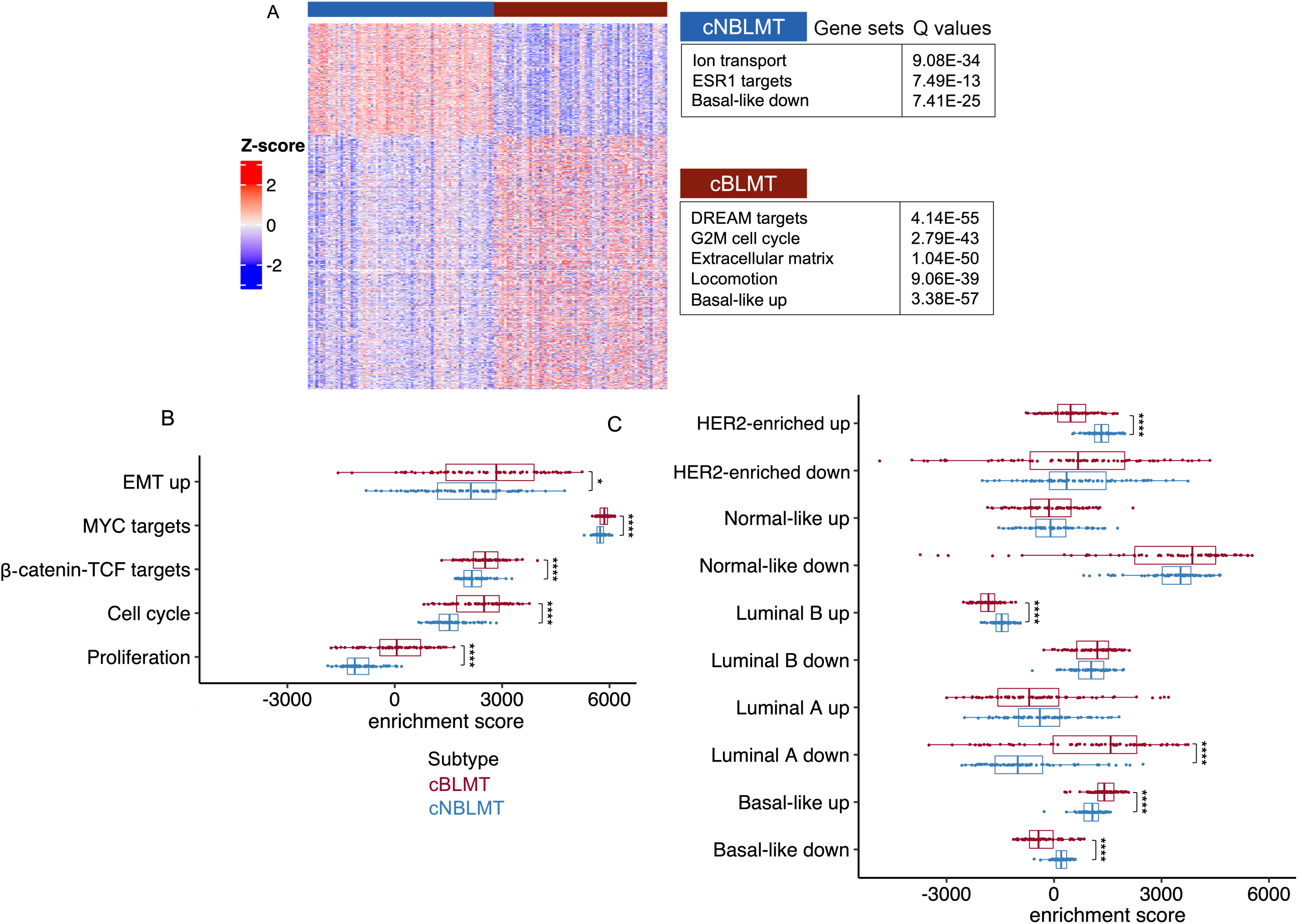
Differentially expressed (DE) gene analysis indicates the enrichment of hBLBC signatures in cBLMT. A. Heatmap of the row scaled log_2_(TPM) values of 1,620 DE genes between cBLMT and cNBLMT samples, identified with an expression fold change of >2 and a BH adjusted p-value of < 0.01 for each DE gene (see Methods). The enriched functions among each DE gene group are indicated. B & C. Distribution of single sample gene set enrichment analysis (ssGSEA) scores of canine tumors with hBLBC signature gene sets (B), as well as with gene sets with expression patterns characterizing each human breast cancer subtype [52] shown (C). P-values are from Wilcoxon tests. *: p < 0.05; ****: p <0.0001.

We then performed single sample gene set enrichment analysis (ssGSEA) with the signature gene sets activated in hBLBC and gene sets known to be up- or downregulated in each human breast cancer subtype [3, 52, 53]. This analysis indicates cell cycle, cell proliferation, MYC targets, β-catenin-TCF targets, and epithelial mesenchymal transition (EMT) genes are all significantly more upregulated in cBLMTs, compared to cNBLMTs (Fig. 3B). Moreover, the gene set known to be highly expressed in hBLBC is significantly upregulated in cBLMTs, while the gene set known to be lowly expressed in hBLBC is significantly downregulated in cBLMTs [52] (Fig. 3C). This pattern is, however, not observed in cNBLMTs, as the gene set known to highly expressed in hLumA tumors [52] does not show significant upregulation in cNBLMTs (Fig. 3C). These results are largely supported by findings with the validation set. The analyses indicate that cBLMT captures key molecular features of hBLBC examined, while cNBLMT fails to do so with those of hLumA.

### The cBLMT subtype contains ER-PR-, ER-PR+, and ER+PR+ tumors

hBLBCs consist of approximately 70% ER-PR- (expressing neither ER nor PR) and 30% ER+PR-tumors, with ER-PR+ and ER+PR+ tumors extremely rare, based on the *ESR1* and *PGR* transcript abundance levels (Figs. 4A-B). However, among cBLMTs, ER-PR+ and ER+PR+ tumors are significantly more frequent, accounting for 33% and 29% respectively, while ER-PR- and ER+PR-tumors only make up 29% and 9% respectively (Figs. 4C-D; Table 1). Notably, about 78% of ER-PR+ cBLMTs were classified as basal-like in PAM50 analysis, similar to the percentage for ER-PR-cBLMT (85%) (Table 1). PAM50 classification also categorized 30% of ER+PR+ cBLMTs as basal-like, a proportion significantly higher than in cNBLMTs (7%) (96% of cNBLMTs are ER+PR+) (Table 1). The observation of higher *PGR* expression in cBLMTs and canine tumors than in human breast tumors (Figs. 4A-D) is supported by other canine and human studies [27, 30, 31, 57]. We noted no significant differences in the dog’s spayed status or the *PIK3CA* mutation status among the ER+/-PR+/-cBLMT subgroups (Figs. 4E-F), indicating that the higher *PGR* expression is unlikely to be associated with either factor.

**Fig. 4.**
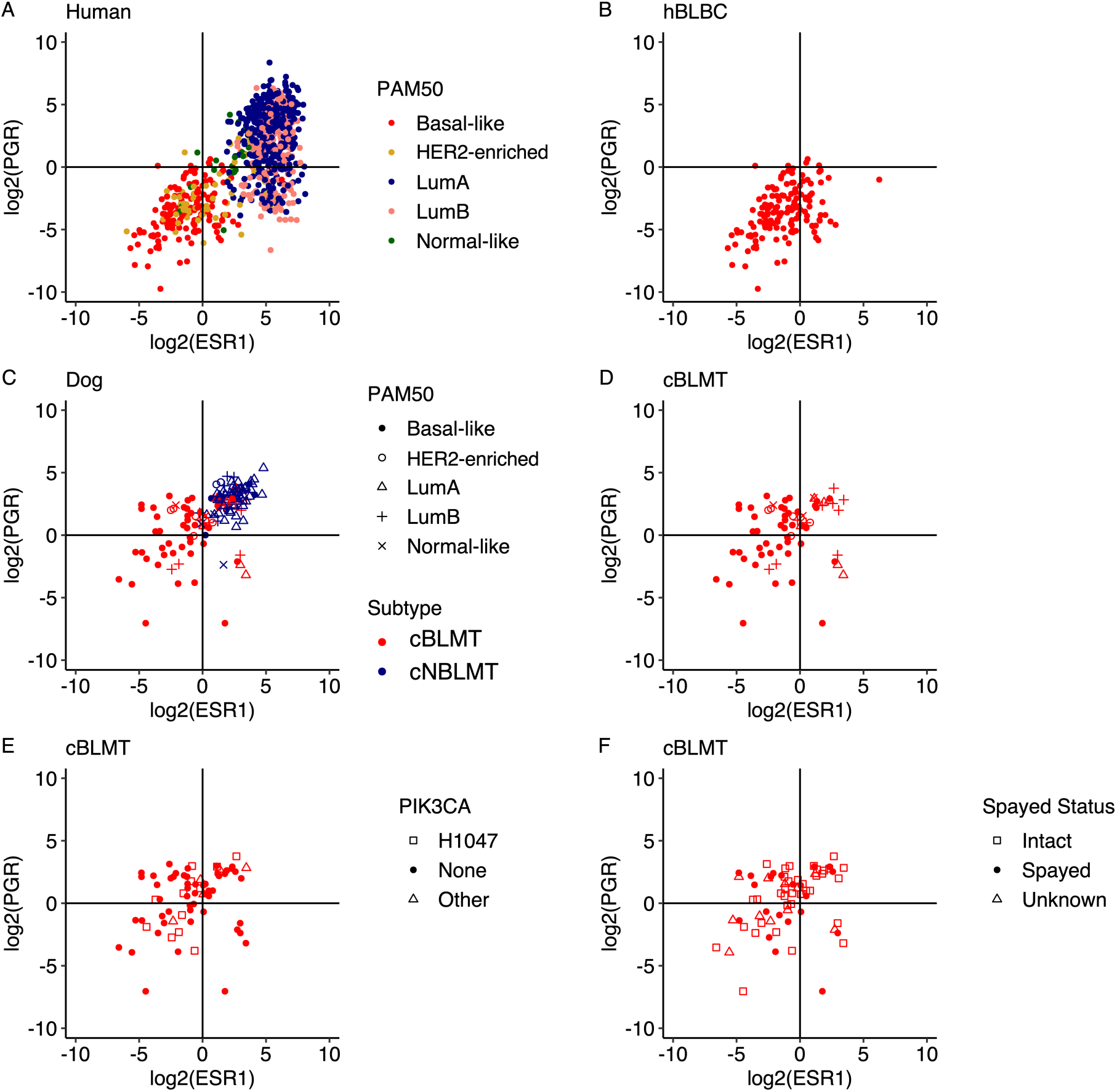
cBLMT contains more ER-PR+ tumors than hBLBC. Scatter plots show the log_2_(FPKM) values of *ESR1* and *PGR* for all TCGA human breast cancers with a PAM50 subtype (n=827) (A), hBLBCs (n=140) (B), both cBLMTs and cNBLMTs (n=143) (C), as well as cBLMTs (n=69) with the PAM50 subtype (D), *PIK3CA* mutation status (E), dog’s spayed status (F) indicated.

### All cBLMT subgroups capture key molecular features of hBLBC

We compared each of ER-PR-, ER-PR+, and ER+PR+ cBLMT subgroups to cNBLMT (the ER+PR-subgroup contains only 6 samples and was thus excluded from the analysis; see Table 1). We found that the DE genes are enriched in functions that characterize hBLBC [3, 7, 58] (Figs. 5A-C; Table 1). Briefly, cell cycle and Wnt signaling are enriched among upregulated genes in each cBLMT subgroup, and the enrichment is especially significant in ER-PR- and ER-PR+ cBLMTs (Figs. 5A-C). Signatures for impaired BRCA2 function and p53 signaling, both known features of hBLBC [3], are also enriched among upregulated genes in ER-PR- and ER-PR+ cBLMTs (Figs. 5A-C; Table 1).

**Fig. 5.**
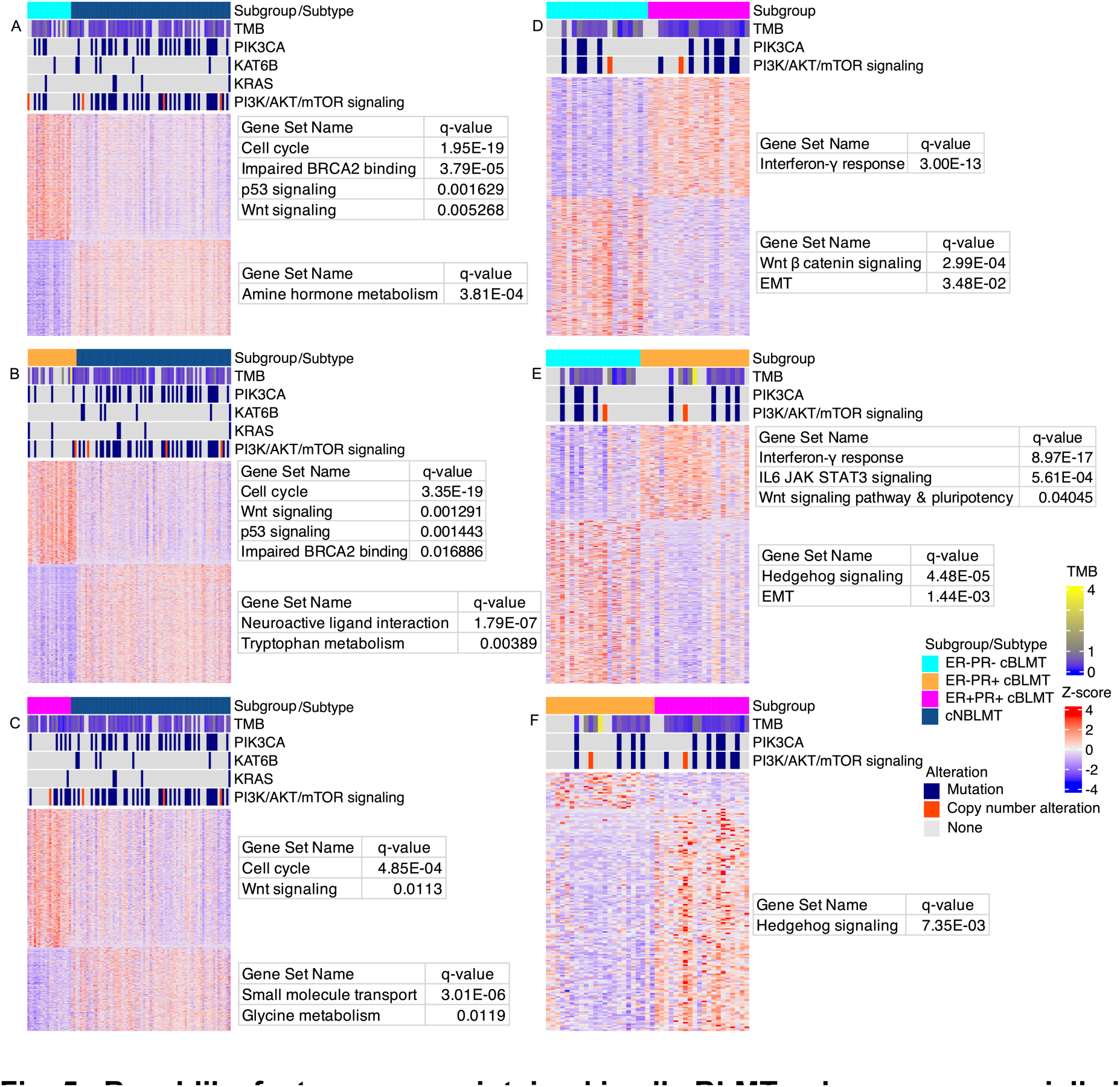
Basal-like features are maintained in all cBLMT subgroups, especially in ER-PR+ and ER-PR-cBLMTs. A-C. Heatmaps showing the DE genes identified between cNBLMT and each of the ER-PR- (A), ER-PR+ (B), and ER+PR+ (C) cBLMT subgroups, with a BH adjusted p-value of < 0.05 and an expression-fold change of > 2. The heatmaps are presented as described in Fig. 3A, along with TMB and the most mutated genes indicated as described in Fig. 1. A tumor was classified ER+ or PR+ if its *ESR1* or *PGR* gene has a FPKM value of > 1, respectively; otherwise, the tumor was classified ER- or PR-. D-F. Heatmaps of DE genes identified between ER+/-PR+/-cBLMT subgroups, presented as described in A-C.

### ER-PR+ cBLMTs harbor upregulated INF-γ response genes but downregulated *IFNG*

We performed DE analysis between cBLMT subgroups and found additional hBLBC characteristics [2, 3, 7, 59, 60] specific to each subgroup (Figs. 5D-F; Table 1). Compared to ER+PR+ and ER-PR+ cBLMTs, upregulated genes in ER-PR-cBLMTs are significantly enriched in functions such as EMT (Figs. 5D-E). Meanwhile, upregulated genes in ER+PR+ and ER-PR+ cBLMTs are enriched in functions including interferon-gamma (INF-γ) response (Figs. 5D-E). Other notable findings include that Wnt-signaling-initiated pluripotency genes and IL6/JAK/STAT3 signaling genes are significantly upregulated in ER-PR+ cBLMTs, compared to ER-PR-cBLMTs (Figs. 5D-E). Hedgehog signaling genes are significantly upregulated in both ER-PR- and ER+PR+ cBLMTs, compared to ER-PR+ cBLMTs (Figs. 5D-E).

Interestingly, while the INF-γ response genes are upregulated in both ER+PR+ and ER-PR+ cBLMTs (Figs. 5D-E), the INF-γ gene (*IFNG*) itself is significantly downregulated in ER-PR+ cBLMTs than in ER+PR+ cBLMTs (Fig. 6A). This indicates that T cell exhaustion may occur in ER-PR+ cBLMTs [59]. To investigate this possibility, we examined the expression of 8 known T cell exhaustion markers [61], but did not find a significant difference among the three subgroups (Fig. 6A).

**Fig. 6.**
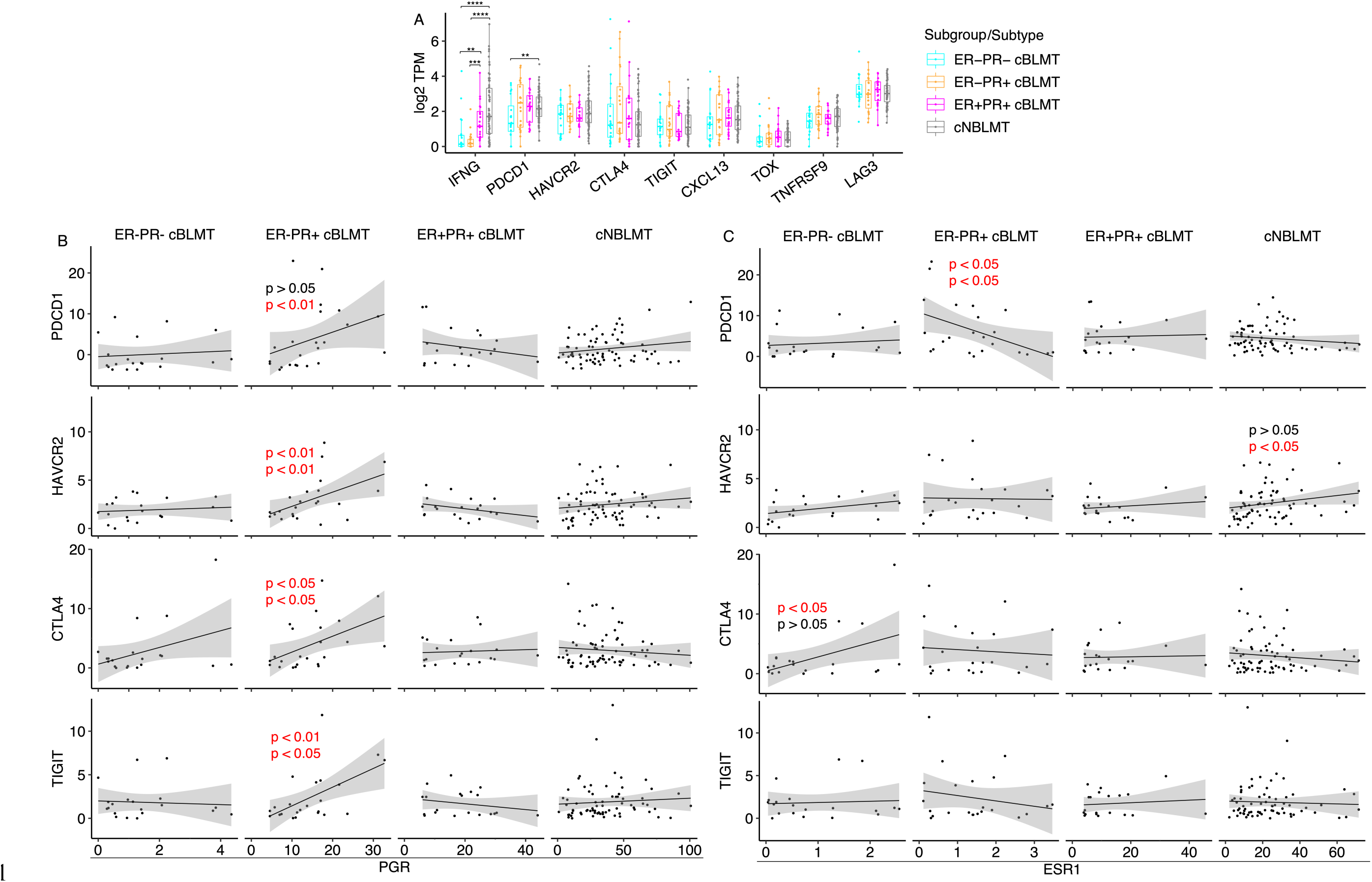
*PGR* correlates with several T cell exhaustion signature genes in mRNA expression in ER-PR+ cBLMTs. A. Dot plots of log_2_(TPM) values of *IFNG* and 8 canonical T cell exhaustion marker genes in each cBLMT subgroup and cNBLMT. P-values are from Wilcoxon tests. **: p<0.01; ***: p<0.001; ****: p<0.0001. B & C. Pearson (top) and Spearman (bottom) correlation analysis between *PGR* (B) or *ESR1* (C) and *PDCD1*, *HAVCR2*, *CTLA4*, or *TIGIT* in mRNA expression in each subgroup and subtype shown. P-values of only significant Pearson and/or Spearman correlations are indicated.

As 6 out of 8 T cell exhaustion markers, including *PDCD1* (encoding PD-1), express higher in ER-PR+ and/or ER+PR+ cBLMTs (Fig. 6A), we examined the association of each marker with *PGR* or *ESR1* in expression. We found that four markers, including *PDCD1*, *HAVCR2*, *CTLA4*, and *TIGIT*, have a significant positive association with *PGR* in ER-PR+ cBLMTs (Fig. 6B). No such associations were found for *PGR* in other cBLMT subgroups or the cNBLMT subtype, or for *ESR1* in any cBLMT subgroup or cNBLMT (Figs. 6C).

### *PGR* expression is associated with gene silencing

To further understand PR in canine tumors, we investigated genes that correlate with *PGR* or *ESR1* in transcript abundance in both human and canine tumors. The same as in hLumA tumors, over 1000 genes were found to be positively correlated with *ESR1* in cNBLMTs (Fig. 7A). Importantly, both sets of genes are enriched in the same functions, including cell cycle (Fig. 7A). For *PGR*, about 2.7 times as many (776 versus 289) positively correlated genes were identified for cNBLMT than hLumA (Fig. 7A). Moreover, while the *PGR*-correlated 289 genes in hLumA tumors are enriched in largely the same functions as those of the *ESR1*-correlated genes, the 776 *PGR*-correlated genes in cNBLMTs are enriched in interferon-γ response (Fig. 7A).

**Fig. 7.**
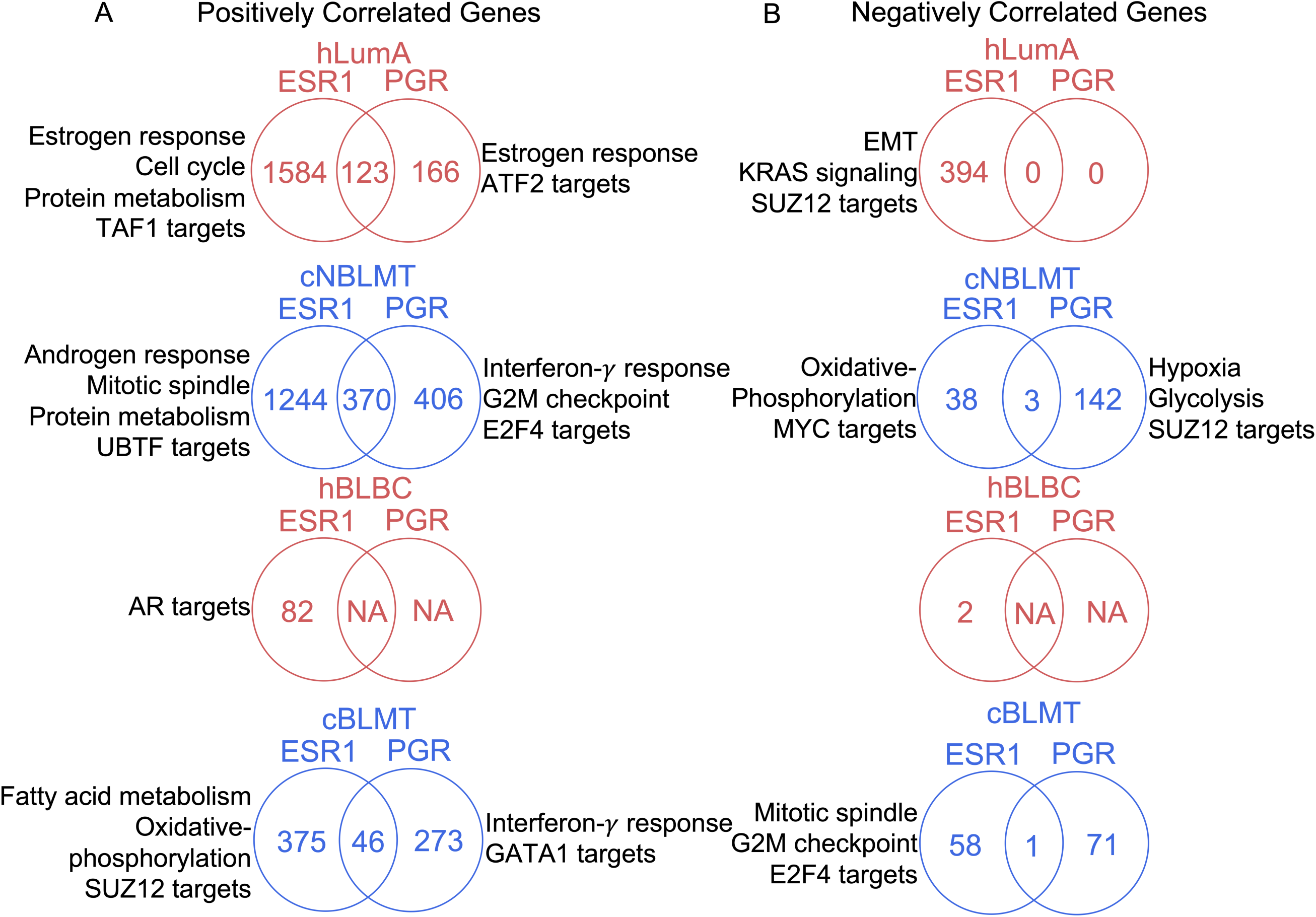
*PGR* is associated with gene silencing in canine tumors but not in human tumors. A. Venn diagrams indicating the number of genes that are positively associated with *ESR1* and/or *PGR* in mRNA expression in each human or canine subtype specified. These genes were identified as those having correlation coefficient 𝑅 ≥ 0.3 and BH-adjusted 𝑝 ≤ 0.05 in both Pearson and Spearman correlation analyses. Indicated are also the top enriched functions of genes that are correlated with only *ESR1* or *PGR*. B. Venn diagrams for genes negatively correlated with *ESR1* and/or *PGR*, identified with 𝑅 ≤ −0.3 and other cutoffs and presented as described in A.

Fewer positively correlated genes with *ESR1* or *PGR* were identified in basal-like tumors in both species. For *ESR1*, we found about 400 genes in cBLMTs, which are enriched in functions including fatty acid metabolisms, oxidative phosphorylation, and SUZ12 targets, and 82 genes in hBLBCs, which are not enriched in any specific functions (Fig. 7A). For *PGR*, while no genes were found in hBLBC, about 300 genes in cBLMT were identified and are enriched in interferon-γ response and GATA1 targets (Fig. 7A).

Negatively correlated genes show a larger dog-human difference, especially for the basal-like subtype and *PGR*. For *ESR1* in non-basal-like subtypes, 394 genes in hLumA tumors, enriched in functions including EMT, KRAS signaling, and SUZ12 targets, and 38 genes in cNBLMTs, enriched in functions such as oxidative phosphorylation, were identified (Fig. 7B). However, in basal-like subtypes, only 2 genes were negatively correlated with *ESR1* in hBLBCs, compared to 58 in cBLMTs that are enriched in cell cycle-related functions such as mitotic spindle and G2M checkpoints (Fig. 7B). For *PGR*, while no genes were identified in either hLumA or hBLBC tumors, 142 genes in cNBLMT, enriched in functions such as hypoxia and glycolysis, and 71 genes in cBLMT, not enriched in any specific functions, were found (Fig. 7B).

## Discussion

Taking advantage of the recently published RNA-seq data for hundreds of canine mammary tumor cases, we performed, to our knowledge, the first genome-wide and independent (not using any known biomarkers) subtyping of this cancer common in bitches that are intact or spayed late. The study identifies two subtypes, and further shows that one subtype molecularly resembles hBLBC, while the other subtype appears not to match any human breast cancer subtypes. This conclusion is consistent with our previous study [14], but differs from a recent publication reporting that canine and human luminal A tumors have more molecular homology than canine and human basal-like tumors [26]. While our canine-only PAM50 analysis also classifies most tumors of one canine subtype as luminal A, the same as the study by Bergholtz et al. [26], our cross-species PAM50 analysis clearly separates canine luminal A tumors from hLumA tumors, unlike basal-like tumors (as such, we named the two canine subtypes as basal-like, cBLMT, and non-basal-like, cNBLMT) (note that Bergholtz et al. [26] did not perform cross-species PAM50 classification). We further show that cBLMTs capture key molecular features and expression patterns of hBLBCs, whereas cNBLMTs fail to do the same with hLumA tumors. Our results are supported by the histology of the canine subtypes. cBLMTs are enriched in simple carcinomas and simple adenomas, where only one cell lineage prominently proliferates. cNBLMTs, however, are enriched in complex or mixed carcinomas and adenomas, where multiple cell lineages (e.g., epithelial cells and myoepithelial cells) proliferate [14, 19, 20]. Complex or mixed tumors (e.g., adenomyoepithelioma) are very rare in human breast cancer [14, 21, 22]; thus cNBLMTs do not match hLumA tumors histologically.

Although cBLMT co-clusters with hBLBC in PAM50 classification and captures the key molecular features and gene expression patterns of hBLBC, cBLMT contains ER-PR+ and ER+PR+ tumors, which are rare in hBLBC. Moreover, while ER+PR-tumors are common in hBLBC and other breast cancer subtypes, ER-PR+ tumors are nearly nonexistent in any human breast cancer subtype. This is because that in human breast cells, PR is induced by ER [62, 63] and thus, without ER, PR will not be expressed. One notable difference between the estrous cycle in bitches and the menstrual cycle in women is the luteal phase, which lasts 14 days for humans but 2 months for dogs. As the canine mammary glands are constantly exposed to a high level of progesterone during the luteal phase [64, 65], it is possible that the *PGR* gene is still actively transcribed after the *ESR1* gene is silenced in dogs, resulting in the ER-PR+ tumors. More studies are needed to investigate this possibility.

Despite the difference in the PR expression status, ER+PR+ and ER-PR+ cBLMTs capture key molecular features of hBLBCs, the same as ER-PR-cBLMTs. These include upregulation of cell cycle genes and Wnt signaling. Upregulation of cell cycle genes could lead to high cell proliferation, a molecular characteristic of hBLBC [3]. Activated Wnt signaling is also a well-known feature of hBLBC [7, 58]. For example, WNT5B, one of the major Wnt signaling molecules, is known to drive the hBLBC phenotype, both in vitro and in vivo, by activating both canonical and non-canonical Wnt signaling [66]. *WNT5B* is upregulated in both ER-PR+ and ER+PR+ cBLMTs. Wnt signaling-driving pluripotency genes are upregulated in ER-PR+ cBLMTs, which likely further drives the basal-like features of these tumors [58].

One notable finding from our study is that in ER-PR+ cBLMTs, interferon-γ response genes are upregulated, but the interferon-γ gene (*IFNG*) itself is downregulated. Moreover, ER-PR+ cBLMTs appear to express more of the immune checkpoint genes *PDCD1* (encoding PD-1) and *CTLA4*, and only in these tumors, *PGR* is positively correlated with *PDCD1* and *CTLA4*. These results are consistent with T cell exhaustion, where T cells are hypofunctional [67]. T cell exhaustion occurs in many human cancers, including hBLBCs [59], and presents challenges and opportunities in cancer immunotherapy [67]. Due to the small sample size, we cannot conclude definitively that T cell exhaustion indeed occurs in cBLMTs. Once this possibility is validated with further studies, cBLMTs could be a valuable model to investigate the relationship among progesterone, PR, and T cell exhaustion. Importantly, cBLMTs may be good models to test novel immunotherapies targeting T cell exhaustion [67].

Our study reveals that unlike *ESR1*, *PGR* is associated with gene silencing in canine mammary tumors. However, the silenced genes appear to be random and not enriched in any particular functions, especially in ER-PR+ cBLMTs. Interestingly, we find that many of the *ESR1* or *PGR*-correlated genes are targets of SUZ12, a component of polycomb repressive complex 2 (PRC2) that primarily methylates lysine 27 of histone H3 (e.g., H3K27me3, a marker of transcriptionally silent chromatin). Further studies are needed to determine if PRC2 is responsible for *PGR*-associated gene silencing.

## Conclusions

We identify two molecular subtypes in spontaneous canine mammary tumors. One subtype, cBLMT, molecularly and histologically resembles hBLBC, a breast cancer subtype that lacks an effective treatment and has the worst clinical outcomes. The other subtype, cNBLMT, appears not to match any human breast cancer subtype molecularly and histologically. While cBLMTs also consist of ER-PR+ and ER+PR+ tumors (which may be related to the long luteal phase of the estrous cycle in dogs), a difference from hBLBC, we note that these tumors capture the key molecular features of hBLBCs, the same as ER-PR-cBLMTs. Thus, cBLMTs could serve as a much-needed spontaneous animal model for hBLBC, filling a critical gap in breast cancer research. Moreover, while much more studies are needed, ER-PR+ cBLMTs may provide a valuable system to study T cell exhaustion, as well as estrogen/ER-independent roles of progesterone and PR in gene silencing.

## Declarations

### Ethics approval and consent to participate

Not applicable.

### Consent for publication

Not applicable.

### Availability of data and materials

The RNA-seq data generated from this study are submitted to the SRA database at https://www.ncbi.nlm.nih.gov/bioproject/PRJNA912710. All other RNA-seq and microarray data sets analyzed in this study are obtained from the SRA and GEO databases at: https://www.ncbi.nlm.nih.gov/bioproject/PRJNA489087/, https://www.ncbi.nlm.nih.gov/bioproject/PRJNA203086/, https://www.ncbi.nlm.nih.gov/bioproject/?term=PRJNA561580, https://www.ncbi.nlm.nih.gov/geo/query/acc.cgi?acc=GSE20718, https://www.ncbi.nlm.nih.gov/geo/query/acc.cgi?acc=GSE22516, and https://www.ncbi.nlm.nih.gov/geo/query/acc.cgi?acc=gse20685.

## Funding

This research was funded by the National Cancer Institute (NCI), National Institute of Health (NIH) grants R01 CA252713 and R01 CA182093. The Ohio State University College of Veterinary Medicine Biospecimen Repository, one of the sample sources, is supported by NCATS UL1TR001070 and NCI P30CA016058.

### Authors’ contributions

SZ, TF, KD, and JW conceived the experiment design. JW and SZ wrote the manuscript. KH performed all QC analysis. YF performed DE analysis and plotted Fig. 3A. KD provided advice on statistical analyses. TF performed initial NMF and PAM50 classification. JW conducted all other analyses.

## List of abbreviations

NMF: non-negative matrix factorization
GEO: Gene Expression Omnibus
NCI: National Cancer Institute
GDC: Genomic Data Commons
TCGA: The Cancer Genome Atlas
cBLMT: canine basal-like mammary tumors
cNBLMT: canine nonbasal-like mammary tumors
ER: Estrogen receptor
PR: Progesterone receptor
TPM: Transcripts per million
FPKM: Fragments per kilobase of exon per million mapped
QC: Quality control
CDS: coding sequence
TMB: tumor mutation burden
hBLBC: human basal-like breast cancer
hLumA: human luminal A breast cancer
EMT: epithelial mesenchymal transition
DE: Differentially expressed
PRC2: Polycomb repressive complex 2

## Acknowledgements

We thank Ms. Jin Qian for her contribution to the study; Dr. Holly Borghese and Dr. William C. Kisseberth for their help in sample collection; the Georgia Advance Computing Resource Center (GACRC) for providing the computing power for this work; Dr. Michael Skaro for advice on figure design; and the authors who published canine and human data used in this work.

